# A sequencer coming of age: *de novo* genome assembly using MinION reads

**DOI:** 10.1101/142711

**Authors:** Carlos de Lannoy, Dick de Ridder, Judith Risse

## Abstract

Nanopore technology provides a novel approach to DNA sequencing that yields long, label-free reads of constant quality. The first commercial implementation of this approach, the MinION, has shown promise in various sequencing applications. The presented literature review gives an up-to-date overview of the MinION’s utility as a *de novo* sequencing device. It is argued that the MinION may allow for portable and affordable *de novo* sequencing of even complex genomes in the near future, despite the currently error-prone nature of its reads. Through continuous updates to the MinION hardware and the development of new assembly pipelines, both sequencing accuracy and assembly quality have already risen rapidly. However, this fast pace of development has also lead to a lack of oversight in the expanding landscape of analysis tools, as performance evaluations are outdated quickly. Now that the MinION is approaching a state of maturity, a thorough comparative benchmarking effort of *de novo* assembly pipelines may be at place.

## 1 Introduction

The development of novel genome sequencing methods has been a major driving force behind the rapid advancements in genomics of the last decades. Notably, the advent of second generation sequencing (SGS) provided researchers with the required throughput and cost-efficiency to sequence many more genomes than was previously deemed feasible. Recent years saw the dawn of what can be perceived as a third generation; one that allows amplification-free reading of single DNA molecules in long consecutive stretches [1, review]. Currently, this new generation is dominated by two methods: nanopore sequencing and single-molecule real time (SMRT) sequencing, championed by Oxford Nanopore Technologies (ONT) and Pacific Biosciences (PacBio) respectively.

Conceptually, nanopore sequencing is easier to explain than most other sequencing methods. An electrical potential is applied over an insulating membrane in which a single small pore is inserted. A DNA strand is pulled through the pore and the sequence is inferred from the characteristic way in which the passing base combinations influence the current. In 1989 David Deamer roughly sketched this concept as it is applied today, although it took more than two decades of key innovations to bring the concept to fruition [2]. Since the introduction of the first commercially available nanopore sequencing device, ONT’s MinION, and the start of the MinION Access program (MAP) in 2014, the field of nanopore sequencing has been advancing at a rapid pace; both new applications and improvements to existing ones are published on a regular basis.

The advantages of the MinION over other sequencing devices are numerous. Both its size, roughly that of a cellphone, and its cost, a thousand dollars for a starter kit, are a mere fraction of that of competitors. Running the MinION is also reasonably time- and cost-effective; an 48-hour sequencing run currently costs around 800 dollars^1^ and yields about 5 Gbases [3, 4]. Furthermore, the technique does not rely on any labeling techniques to recognize different bases, while Sanger, SGS methods and SMRT do require some form of labeling of nucleotides. Amplification by PCR is optional for the MinION, while this step is mandatory for Sanger and SGS-methods. Not only does omitting these steps simplify sample preparation for MinION samples; it also helps to avoid errors and biases (e.g. the CG-bias for PCR) and allows detection of modified bases [5]. Finally, the maximum read length produced by the MinION is many times greater than that of both second-generation and Sanger sequencing and only paralleled by SMRT sequencing, which is highly advantageous in resolving repeat sequences.

The most prominent disadvantages of the MinION with respect to its competitors are the lower signal-to-noise ratio, stochasticity introduced by its biological components and the resulting high error rate of basecalling. Indeed, the MinION is a product in development and the used materials (i.e. membranes, nanopores and buffers) are still being optimized. Furthermore, it is thought that significant improvements are still possible in the software pipelines that translate current signal to DNA sequence, even though recent efforts have increased the base-calling accuracy dramatically, from around 85% in the first half of 2016 [6–8] to over 95% in March of 2017 [3].

In this literature review, an up-to-date overview of several aspects of nanopore sequencing is given. First, the physical sequencing process as it takes place inside the MinION is outlined. Then, the general structure of analysis pipelines is described, along with currently available software implemented in these pipelines and their respective strengths and weaknesses. It should be noted that nanopore sequencing is a rapidly advancing field. While some work discussed in this chapter is considered cutting-edge at the moment of writing, the reader is advised to keep the publication date of said work in mind.

## 2 Physical basis of DNA sequencing using nanopores

In a nutshell, the underlying principle of nanopore sequencing can be explained as follows: a microscopic opening wide enough to allow single-stranded DNA to pass - the nanopore - is introduced in an insulating membrane between two compartments filled with saline solution and an electric potential is applied over it. DNA strands are then added to one compartment and allowed to diffuse toward the nanopore, where they are captured by the electric field and threaded through the pore. While a strand is passed through, the characteristic way in which the bases influence the electric current through the nanopore is measured. These measurements can then be decoded to retrieve the sequence of the DNA strand (Figure 1).

**Figure 1:**
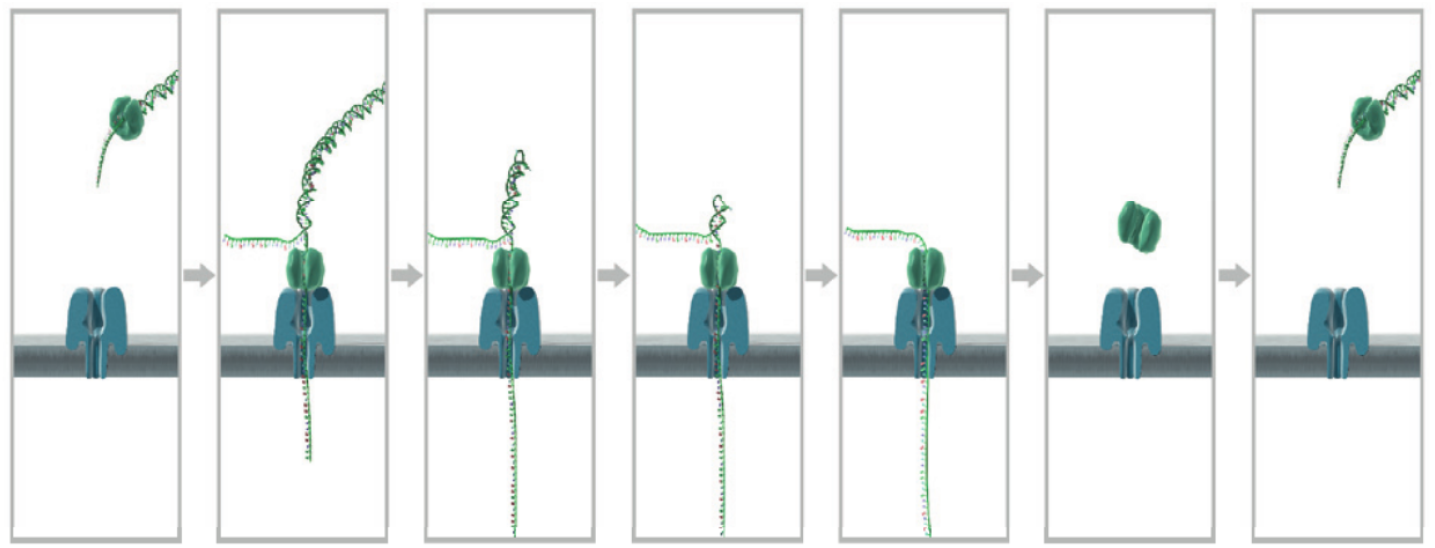
Sequencing of a DNA strand using nanopores. From left to right, double-stranded DNA with attached motor protein attaches to a pore protein in an insulating membrane. The applied potential pulls one strand through the pore, while the motor protein unzips the DNA in a step-wise fashion. After the DNA has been unzipped completely and one strand has passed through, the complex detaches from the pore entrance and the pore is ready to receive another strand. Taken from [6].

In recent years, several key discoveries have brought reliable nanopore sequencing closer at a rapid pace. In a step-by-step exploration of the sequencing process, these discoveries will be discussed next.

### Choice of pore: biological versus solid-state

Nanopore sequencing efforts are sub-categorized in two groups based on the choice of nanopore. Most current efforts implement biological nanopores, which are protein polymers derived from naturally occurring counterparts. Through genetic engineering, biological nanopores are modifiable in terms of dimensions and placement of electrical charge. These properties are also highly reproducible from one pore to the next. Functionality can be further modified by attaching compatible enzymes to the pore opening. Like their naturally occurring counterparts however, they need to be embedded in lipid bilayers, which are generally prone to disruption, particularly when exposed to varying electrical potentials. In the MinION, this was partly solved by constructing membranes out of a more stable single layer of polymers, rather than the traditional bilayer. Solid-state nanopores on the other hand are made by burning openings in a synthetic membrane using a focused electron or ion beam [9]. Contrary to biological nanopores, solid-state nanopores are compatible with a wide range of strong and chemically stable materials with equally diverse properties. Pores are also more easily parallelized and integrated in electrical readout circuits. A major disadvantage at the moment is the reproducibility of the pore dimensions. They also do not combine as easily with modifying enzymes. As a result, solid-state nanopores currently produce noisier and less easily interpretable signals than biological nanopores. In the following, the focus will lie on biological nanopore sequencing and the term nanopore will refer to the biological kind.

### Structure and charge of the nanopore

One important structural property that makes a biological pore suitable for DNA sequencing is a constriction site at which the passing strand exerts the most influence on the electrical current. The length of the constricting passage largely determines how many bases simultaneously influence the electrical current and thus the number of bases *k* that is “read” simultaneously at a given time. *k* should be kept low enough to allow recognition of a signature current for each different combination of *k* bases, or *k*-mer, and high enough to allow for some overlap between subsequent *k*-mers, as this benefits basecalling accuracy by allowing every base to be read *k* times. Modified versions of both pore proteins that have seen application in the MinION, MspA (designated R7 by ONT) and the currently used CsgG [10] (designated R9, Figure 2), have a constricted passage that allows detection of manageable *k*-mer lengths. for the 10Å-long constriction of the CsgG pore, basecalling models assume that five nucleotides influence the current at one given time sufficiently to discern all different nucleotide combinations and thus 5-mers are called.

**Figure 2:**
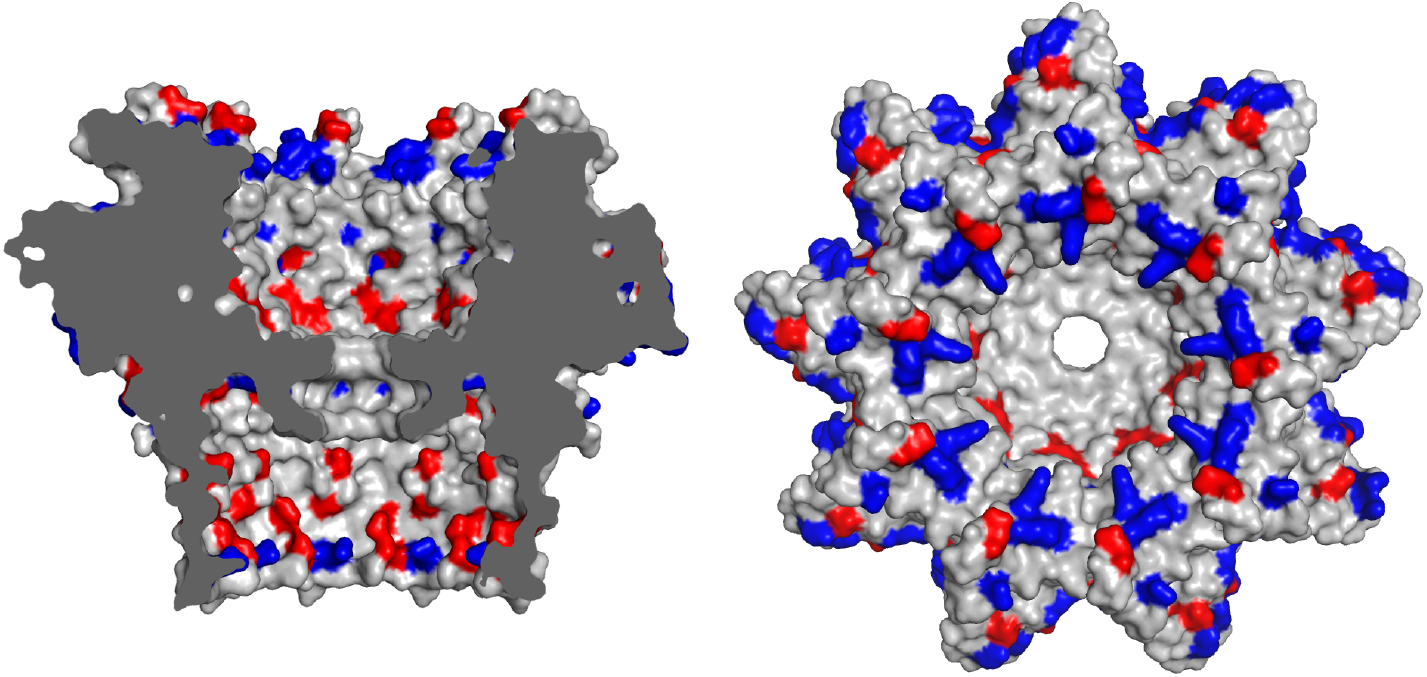
Protein structure of the CsgG pore protein complex, a variant of which (R9) is used in current generation MinION flow cells. Positive and negative residues are colored blue and red respectively. PDB ID: 4UV3 [10].

For sequencing to commence, a DNA strand first needs to diffuse towards one side of the pore, referred to as the cis-side, where it is captured by the electric field resulting from the applied potential. It is then threaded through the pore and extruded at the other end, called the trans-side.

Two forces should be considered. First and most importantly, the electrophoretic force induced by a positive electric potential applied at the trans-side attracts the negatively charged DNA and pulls it in. As negative particles leave the cis-side and positive particles simultaneously move in the opposite direction, a positively charged zone forms around the cis entrance of the pore, strengthening attraction of DNA strands. Secondly, strand translocation is influenced by the electro-osmotic flow (EOF), the force induced by the net water and ion flow through the pore. While a DNA strand is in the pore, the EOF normally opposes the direction of the electrophoretic force and thus of translocation, however this effect is relatively minor.

Through iterative optimization of internal architecture, it was found that positive internal surface charges are important for efficient DNA capture [11, 12], while base recognition was found to improve with bulky or hydrophobic amino acid side chains placed at the constriction site as these direct ion flow toward the DNA strand [13]. Although the structure of the modified CsgG pore [10] implemented in the current MinION flow cells (R9-type) has not been publicly released by ONT, modifications to these properties have likely been made.

### Processive control

It should be noted that the processive speed of the strand is too high without any further modifications (between 2·10^6^ and 10·10^6^ bases/s in wild-type MspA) [11]. Currently, the most successful way to exert control over the speed has proven to be the addition of a motor enzyme, phi29 DNA polymerase (DNAP) [14]. In a preparatory step, a poly-T or “leader” adapter is attached to the double-stranded DNA. The DNAP attaches to this adapter, but due to specialized bases in the adapter sequence (left unspecified by ONT), cannot unzip it at this stage [15]. Once the complex attaches to the cis-side of the pore, the blockade is released, presumably due to the force exerted on the strand as demonstrated by [16]. The DNA is fed through the pore in a step-wise fashion (but due to absence of divalent cations and dNTPs, without actually copying the strand like it would normally do [17]), where it can now be “read” at a reasonably regular pace. A modified phi29 DNAP is used in the MinION. The latest release of this motor protein at the moment of writing (dubbed E8) maintains an average throughput speed of 450 bases/s [18].

### Reading the DNA strand

During a MinION sequencing run, the potential over the membrane is kept stable while the electrical current (in the pA-range) is sampled at a frequency in the kHz range (Figure 3). This signal is characteristic for the subsequent bases moving through the pore and will ultimately serve as the basis for basecalling. As the amount of electrolyte is increasingly depleted during the run, the applied potential (typically starting at −180mV) is further decreased by 5 mV per two hours of runtime and increased by 5mV when the MinION switches to another set of wells filled with fresher buffer (see next section).

**Figure 3:**
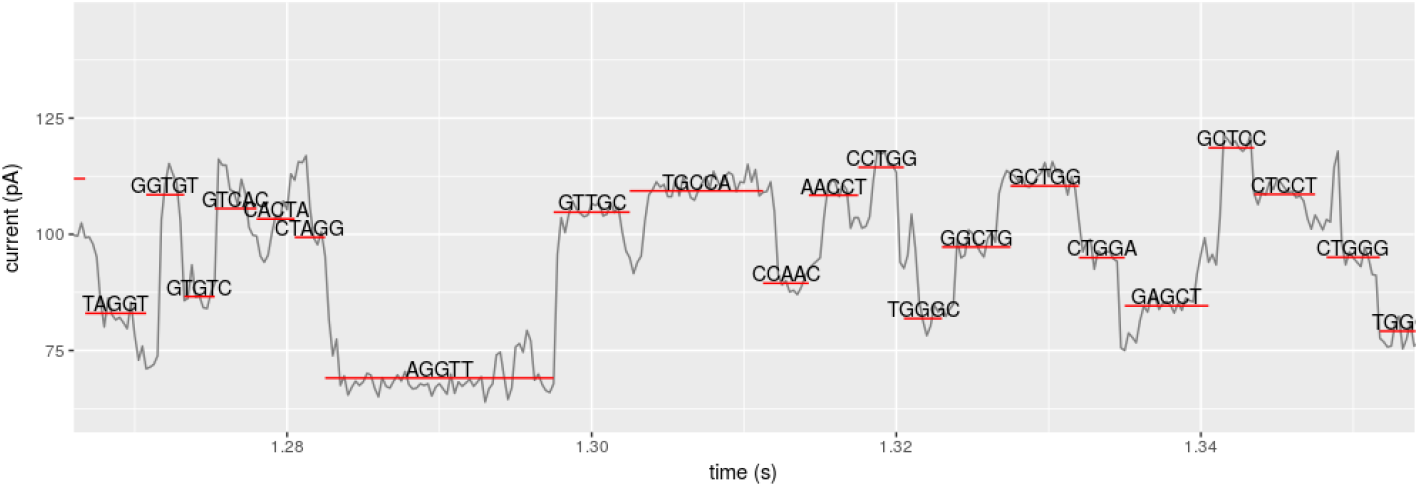
Example of a MinION DNA read as raw data (grey line) and the event data (red lines) extracted from it, corresponding to discrete sets of bases.

While the MinION can read the first strand of a dsDNA-stretch that is threaded through the pore - by definition, the template strand - and discard the complementary strand, it is possible to instead read the complementary strand immediately after the template, thus performing a second read of the same stretch (Figure 4). Combining reads of both strands has been shown to increase sequencing accuracy significantly [6]. So far, ONT has provided two ways of doing so; 2D-sequencing and its successor 1*D*^2^-sequencing (versus 1D sequencing if only the template strand is read). To enable 2D-reads, an abasic hairpin sequence (i.e. a bare backbone of DNA) is attached to the dsDNA, connecting the 3’-end of the template and the 5’-end of its complement. After the template strand has been read, the hairpin structure is threaded through te pore, recognizable due to its abasic nature, followed by the complement strand. The mirroring reads are then decoded jointly so that any sequencing errors may be corrected. It should be noted however, that decoding 2D-reads currently comes at a high computational cost [21]. Furthermore, the exact start and end of the hairpin is not always clear from the signal. The hairpin may also ligate strands in other ways than the intended one; for example, multiple reads may be linked together by hairpins into chimeric reads [22]. Lastly, the read of the complement strand is often threaded through the pore at lower speed [6], possibly due to the force enacted by the strand rezipping while the complement strand is still passing through the pore. In May of 2017, 1*D*^2^-read chemistry was introduced as a replacement of 2D-read chemistry [20]. 1*D*^2^ chemistry does not join the two strands, but rather attaches an adapter to the complement strand, allowing it to attach to the membrane while the template strand is read. Shortly after the template strand has completely left the pore, the complement strand is pulled in and sequenced. Without the hairpin connecting the strands, recognition of its start and end is no longer required and, as the strands are effectively read as two separate reads, no drop in read quality is observed for the complement strand.

**Figure 4:**
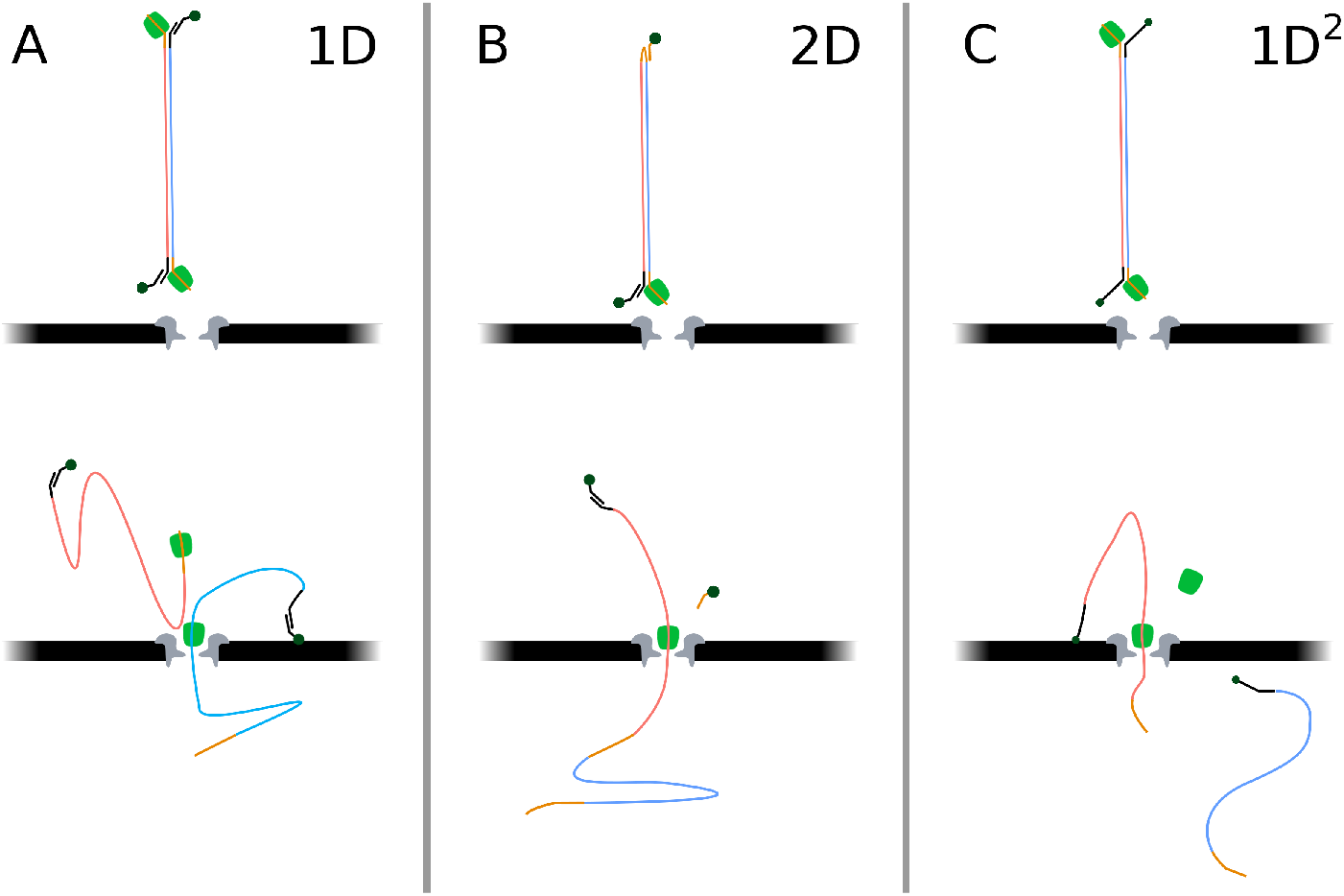
The three categories of DNA reading chemistries for the MinION. (**A**) When using 1D chemistry, only the template strand (blue) is threaded by the motor protein (green) and read. The complement strand (red) is discarded at the cis side of the pore. The tethers (dark-green) allow for selection of properly ligated complexes during sample preparation and attach to the membrane to increase the availability of strands near pores during sequencing. (**B**) The now-deprecated 2D chemistry connected template and complement strand using a hairpin, thus allowing sequencing of the complement strand immediately after the template strand. (**C**) 1*D*^2^ chemistry, the successor of 2D, also allows sequencing of both strands, but rather than attaching the two, the complement strand is tethered to the membrane while the template is sequenced. After the template strand is threaded through, the complement strand is drawn in and the tether is pulled loose. Based on [19], [20] and [15].

### Channel parallelization

Lastly, throughput can be greatly increased by reading the signal from multiple pores in parallel. The current generation of the MinION’s disposable cartridges, called flow cells, can read the signal of up to 512 pores in parallel (Figure 5). The flow cell is equipped with 2048 wells, which are connected in groups of four to multiplexers (MUXs), the switches that control which of the four cells per group is controlled and read out by the circuits. During the initial platform quality check, DNA strands (of unreleased source and sequence) present in the buffer with which the flow cells are shipped is sequenced to discern wells containing precisely one correctly inserted pore from wells in which correct pore insertion has failed (e.g. no or multiple pores were inserted) [23]. The latter scenario may occur, as the insertion of pores is a stochastic process. In a second quality check, the MUX scan, each MUX chooses three wells in order of signal quality and begins readout in the best-quality well. As well quality is expected to decline during the run, the standard protocol switches to the second-best quality pore after eight hours, and the third-best quality after another eight hours. This way, the best and most output is expected in the first part of the run. While a run using a group of wells is in progress, the circuits connected to the MUXs regulate the current in each selected well individually. This also allows expelling of eventual blockades from a pore, by temporarily reversing the current in the affected well while the rest of the wells continue to function normally.

**Figure 5:**
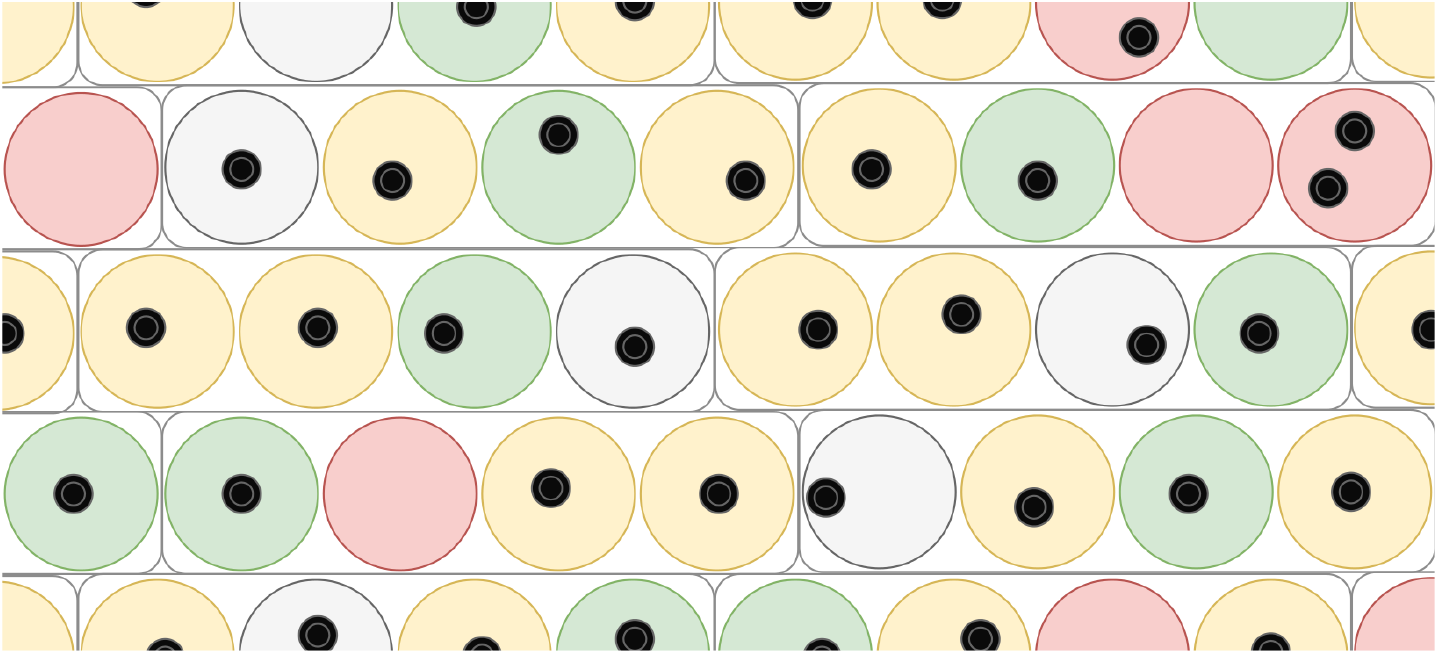
Layout of a MinION flowcell grid. Large circles denote wells in the grid, small black circles denote inserted nanopores. Each group of four wells is controlled by a multiplexer (MUX). During an initial quality check, wells that are unusable due to erroneous pore insertion are removed (red circles). Right before sequencing, the wells are tested a second time and three wells per MUX are ranked on signal quality (if possible). Sequencing of the sample will then commence, starting read-out from the best-performing well (green) and switching to second and third best (yellow) after eight hours.

## 3 Currently available software for MinION sequencing data

Following the process in section 2, a current signal is obtained which needs to be translated into the corresponding DNA sequence. Pipelines used for this purpose generally subdivide the signal into discrete stretches corresponding to a particular set of *k* bases, called events, and then attempt to find the most likely set of bases for each event [7, 8, 24]. Ideally, the processive control exerted by the motor protein causes the strand to move one base further through the pore between each event transition, such that each base in the *k*-mer is read *k* times. Due to irregularity in the motor protein functioning and noise in the signal, this is not always the case. Three transition scenarios may be observed. (I) If the strand progresses exactly one base further from one event to the next, this is called a “step”. Alternatively, (II) the next event may denote a *k*-mer located one or multiple bases further upstream, which is referred to as a “skip”, or (III) the segmentation process may incorrectly split up an event in two events that correspond to the same *k*-mer, which is called a “stay”. Steps, skips and stays may be identified by comparing the bases in the read *k*-mers, however this proves more difficult in short tandem repeat regions as subsequent events would look the same for multiple transition scenarios. More specifically, in a homopolymer stretch of length *k* or longer the difference between skips, steps and stays may be unclear. Similarly, repeats of *m* bases may cause problems in discerning skips of (multiples of) *m* bases and stays. Such ambiguities have been shown to be a major cause of deletions in current basecallers [7, 19, 25, 26].

To assess the quality of basecalling performance, a 3.6 kbase calibration strand derived from the Lambda genome may be added to the sample [15]. The software controlling the MinION (MinKNOW) automatically detects reads derived from the Lambda genome and separates those from the sample reads. Some software tools use these strands to parameterize basecall correction alogorithms [27, e.g.].

Once reads have been basecalled, they may serve several purposes. If SGS reads are available, then those may then be mapped to the MinION reads to correct sequencing errors pre-assembly [28], or to create large low-error contigs. The latter goal may be achieved either by using MinION reads as scaffolds to properly align short reads [29, 30] or by creating short accurate seed regions from short reads, the gaps between which are then bridged by MinION reads [26]. Both approaches were shown to result in accurate and highly continuous *de novo* assemblies and identification of repeats that were collapsed in SGS-only assemblies [26, 31]. In MinION-only assembly pipelines, overlaps between the basecalled sequences are sought to create initial assemblies. Currently, the error-prone nature of MinION reads necessitates a post-assembly error correction (“polishing”), in which raw reads are mapped again to the assemblies until a better consensus is reached [32]. Below, the different tools MinION users have at their disposal to fill in the steps in this pipeline are discussed, along with their strengths and weaknesses.

It should be noted that the presented list of tools is not exhaustive; the focus of this section lies on tools that can be used in *de novo* MinION sequencing, without the need for other sequencing methods. Furthermore, only tools that provide a full solution to their respective step in the assembly pipeline are reported here. Notably, stand-alone read mapping tools exist which may be used in combination with the assembly tools described in this section, however detailed descriptions are omitted for the sake of brevity. Readers with an interest in these alternatives may want to look into GMAP [33], Graphmap [34] and MHAP [35]. As current assemblers either include their own error correction module [36] or work with uncorrected reads [26, 37–39], stand-alone pre-assembly error correction tools are excluded as well.

### 3.1 Segmentation and event detection

Before basecalling takes place, current basecalling tools require the continuous signal to be divided into discrete sections, called events, each supposedly representing one combination of bases. Furthermore, the poly-T lead preceding the sequence needs to be removed and, if 2D chemistry is used, the signal derived from the template strand needs to be separated from that of the hairpin and the complement strand. The latter process is commonly referred to as segmentation. The order in which segmentation and event detection is executed may differ. For example, for ONT’s own analysis pipelines it is noted that segmentation is performed first when basecalling occurs locally, while event detection took place first if the (now discontinued) cloud-based EPI2ME platform service was used [24].

For locally basecalled reads, both segmentation and event detection are performed by the MinKNOW software provided by ONT. The software identifies the hairpin by a characteristic double current spike, while a long poly-T signal denotes the lead. For event detection, MinKNOW was reported to calculate a simple t-statistic between sliding adjacent windows of set size [21]. Peaks in the t-statistic above a certain threshold are then assumed to signify the border between two segments. Optionally, the raw current signal can also be reported for further analysis.

### 3.2 Basecallers

Several dedicated basecalling tools are already available to MinION users. In this section, the underlying principles and implementation of these tools are explored, along with their reported subsequent strengths and weaknesses. It should be noted that an all-inclusive fair comparison, in which all basecallers are optimized for the latest MinION chemistry and then run on the same dataset, is currently lacking. Any comparison in this chapter is therefore based on accuracy reports made by the authors of the open-source basecallers in their publications (i.e. between the Metrichor basecallers and their own).

#### Metrichor basecallers

Metrichor, a spin-off of ONT and its main developer of proprietary analysis software, maintains a range of basecallers that have remained the go-to option for most MinION users. Initially, the Metrichor base-callers relied on hidden Markov models (HMM) to find the biological sequence underlying the segmented signal. As of early 2016, this model was replaced by a more accurate recurrent neural network (RNN)-implementation. Currently several RNN-based versions exist under different names; Albacore, Nanonet and the MinKNOW integrated basecaller. Albacore and the MinKNOW version are stable versions intended for regular MinION users. As of version 1.0.1, Albacore was reported to improve homopolymer calling through the implementation of a transducer, a feature that was transferred from the Scrappie basecaller (see below) [40]. Nanonet is an unsupported version under continuous development and makes use of the CURENNT library to implement its RNN [41]. A cloud-based version was previously integrated in the EPI2ME platform, but this service has been discontinued.

Due to the proprietary nature of the software, the source code of Albacore and the MinKNOW basecaller is currently only open to developer users, while that of Nanonet is openly available [42]. The community manual describes the general procedure followed by all Metrichor basecallers as follows [43]. Long reads are broken up into partially overlapping stretches, referred to as chunks. This keeps the Viterbi-decoding process, required after the RNN has determined *k*-mer probabilities, tractable. The RNN then assigns a likelihood for every possible five-mer to each event in the chunks.

Metrichor basecallers were reported to use a bidirectional long-short term memory (BLSTM)-RNN [21] (Figure 7A). To calculate a hidden state’s output, or activation, a regular (i.e. “vanilla”) RNN takes a linear combination of the activation of the previous hidden state and the current input and then non-linearly transforms it to a value between −1 and +1 (often using a hyperbolic tangent function). A BLSTM-RNN uses a more elaborate structure, the long short-term memory (LSTM) unit [44], which keeps an additional memory composed of previously processed signals and modulates the way in which the memory is updated, forgotten and used in calculating the current activation. A BLSTM-RNN has several hidden layers of LSTM units, some of which use the sequence of signals from one direction to form their memory and some from the opposite direction.

**Figure 6:**
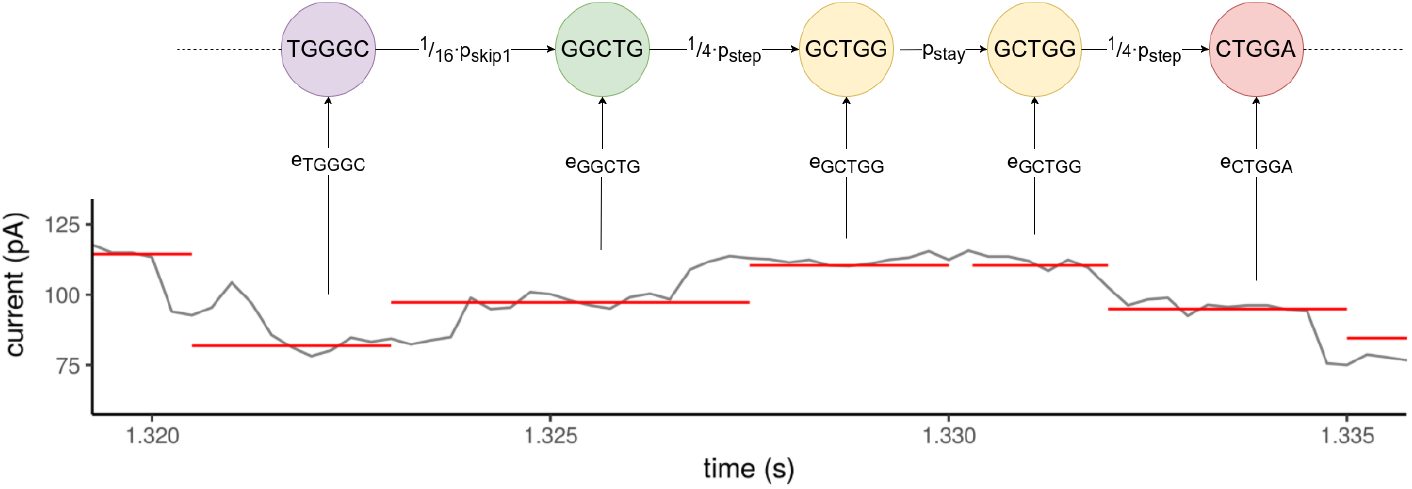
Example of the most likely path of *k*-mers, derived by Nanocall from event data by a hidden Markov model (HMM). The path is chosen by maximizing the product of transition probabilities *p_stay_, p_step_* and *p*_*skip*1_ (modified by fractions in the case of skips and steps, as there are multiple states to which these transitions could lead) and emission probabilities of *k*-mers.

**Figure 7:**
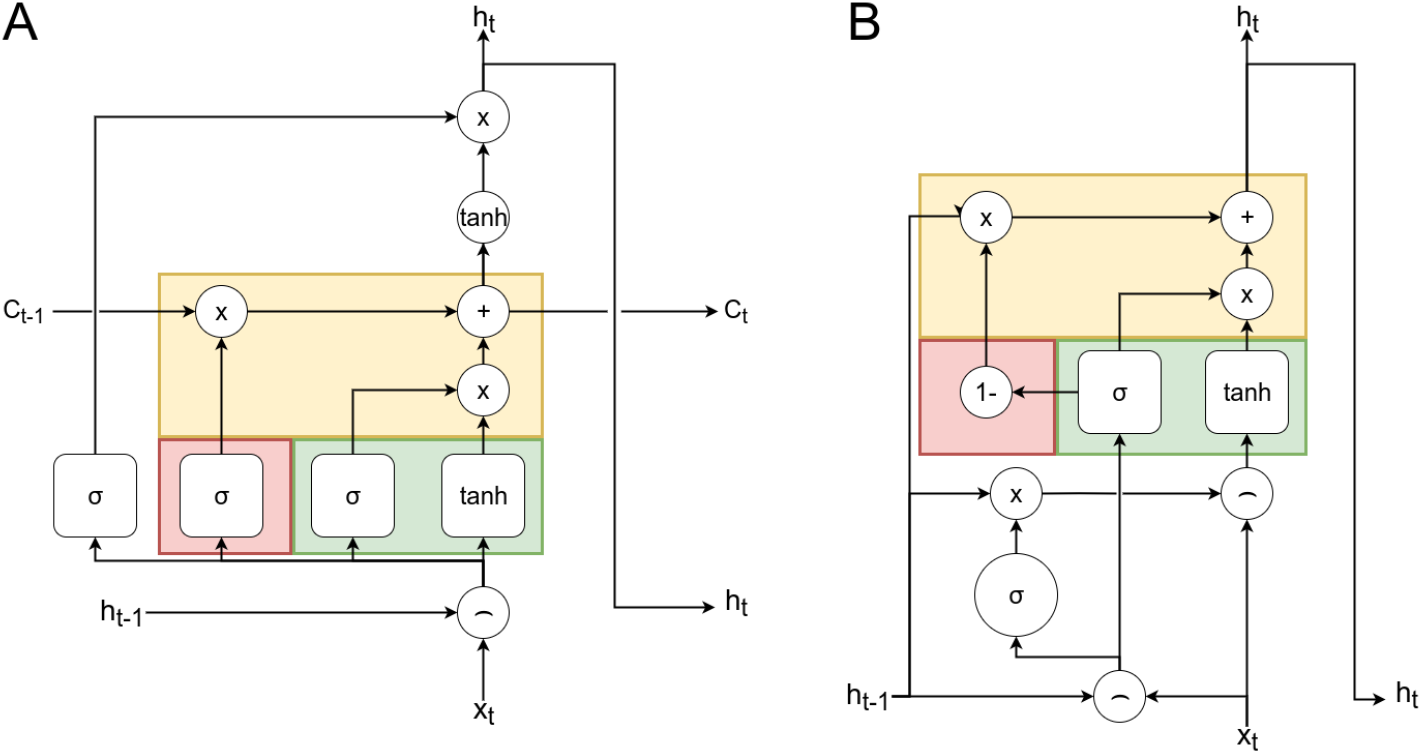
Schematic representation of two types of memory cells used in RNNs. Each requires a set of pointwise operations (circles) and neural network layers (rounded squares) to calculate the activation of the next hidden layer. Colored squares denote named substructures: the forget gate (red), input gate (green) and output gate (yellow). (**A**) The long short-term memory (LSTM) unit maintains a separate memory stream (*C_t_*) which allows information to be passed on updated but unmodified otherwise. (**B**) The gated recurrent unit (GRU) is simpler in design and does not require a separate memory stream. Note that the forget gate is no longer a neural network layer. Forget and input gates are considered fused into a so-called reset or update gate, however for the sake of comparison elements have been depicted in the same structure as the LSTM. Adapted from [52].

RNNs implementing LSTM units have two advantages over vanilla RNNs. First, LSTM units are able to use more information of the previous steps to determine activations in a broader context (in the case of a basecaller, multiple adjoining *k*-mers). Second, as depth and size of the network increases, vanilla RNNs become more difficult to train using back propagation than BLSTM-RNNs, as any updates to the weights of subsequent nodes are reduced too quickly in the process. Thus, layers closer to the input nodes are trained much slower, an issue commonly referred to as the vanishing gradient problem. In LSTM units however, the memory also serves as a shortcut to propagate the signal free of modification of the weights and thus of vanishing gradients. As a result, BLSTM-RNNs do not suffer from the vanishing gradient problem [45].

A softmax function transforms the outputs of the last hidden layer into a posterior probability per five-mer for each event. Posterior probabilities are used to estimate basecalling quality scores (i.e. q-scores). Five-mers that would entail a forbidden move considering the likelihood of surrounding *k*-mers are removed at this point. The maximum likelihood sequence is then extracted from the five-mer likelihoods through Viterbi decoding. Finally, basecalled chunks are merged again.

The accuracy of the Metrichor basecallers is currently considered to be the highest of all available basecallers. Before the switch to RNNs, an international performance analysis estimated the accuracy at 88% for 2D basecalling [6]. David *et al*. reported an accuracy of approximately 68% [7] for 1D basecalling on human data, while Boza *et al*. reported 77% for 1D and 87% for 2D on *E. coli* and *K. pneumoniae* genomes. After moving to an RNN-implementation (as well as numerous improvements in used chemistry and hardware) this percentage has risen to over 95% [3].

#### Scrappie

A proprietary basecaller by Metrichor and platform for ongoing development, Scrappie was the first basecaller reported to specifically address homopolymer basecalling. In previous MinION sequencing efforts, correctly determining the length of homopolymers proved challenging, as the separation between states in a homopolymer stretch was not clear [26, 27, 46]. Scrappie has been reported to process the raw current signal, rather than the event-detected data.

As an initial performance assessment, an early-development version of Scrappie was run in parallel to other Metrichor basecallers on reads of the human chromosome 20. It was found that Scrappie indeed called homopolymer stretches more accurately than the EPI2ME basecaller and Nanonet; homopolymeric stretches of up to 16 bases were called accurately (although a slight over estimation of repeat units was seen between 5 and 15 bases), after which the accuracy fell. In contrast, the EPI2ME basecaller and Nanonet failed to call stretches longer than five bases. The effect was also shown in an analysis of 5-mer counts; Of the tested basecallers, Scrappie stayed closest to the reference genome and did not underrepresent homopolymeric 5-mers [4]. Although these results are promising, a thorough assessment of Scrappie in a later stage of development is required to evaluate the true potential of the transducer-based homopolymer calling.

#### Nanocall

Nanocall [7] was the first open-source basecaller for the MinION offered as an alternative to the proprietary Metrichor software. It was written in C++. Nanocall accepts the segmented signal from minKNOW and assigns *k*-mers to the events using a hidden Markov model (HMM).

An HMM assumes that, in a sequence of events, the probability of the next event being of a certain nature - its state - is dependent only on the current state. This makes sense for the basecalling of subsequent *k*-mers, as the prefix of the next *k*-mer that is called should be the suffix of the previous. The “hidden” property of HMMs refers to the fact that the sequence of states - the path - is not known. Instead, it has to be inferred with some probability from a derived signal. In the case of a basecaller, this signal consists of electrical current levels which are derived of a sequence of *k*-mers. The goal of the HMM is to find the path that is most likely to have emitted the given signal (for a more thorough introduction to HMMs, see [47]).

As described by David *et al*. [7], the HMM underlying Nanocall takes stay, step and skip scenarios into account at each event transition. Although skips of more than one base may occur in a sequence, Nanocall only considers one-base skips to cut down on computational requirements, thus reducing the number of neighboring states to 21 (i.e. stay, four different steps and sixteen different one-base skips). During an initial training phase, state transition probabilities for stays and skips of any number of bases, *p_stay_* and *p_skip_* respectively, are estimated using the Baum-Welch algorithm (*p_step_* is derived as 1 − *p_stay_* − *p_skip_*) [48]. The transition probability for exactly one base *p*_*skip*1_ is then derived as *p*_*skip*1_ = *p*_*skip*1_/(1 + *p_skip_*). Note that all sixteen one-base skips have the same transition probability (i.e. 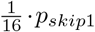), as do the four step scenarios (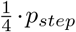). To calculate emission probabilities, Nanocall relies on ONT’s pore models, which provide mean *μ_k_* and standard deviation *σ_k_* of the Gaussian distribution 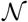 modeling the current level that is characteristic for each different *k*-mer event, as well as mean *η_k_* and standard deviation *γ_k_* of the inverse Gaussian *IG* modeling each current level standard deviation. However, these parameters require fine-tuning to the individual pore and the time passed since the strand entered the pore *i*, as pore characteristics and buffer properties vary from one pore to the next and change during use. To that end several additional parameters are introduced, which are estimated separately for each read using expectation maximization. A linear model adapts the mean to the individual pore for every k-mer using the parameters *shift* and *scale*, while *drift* adjusts it for the time passed since the strand entered the pore. *var* and *var′* scale *σ_k_* and *γ_k_* respectively. Concluding, the emission probability of *k*-mer *k* for event *i* is estimated by multiplying *mean_i_* and *stdv_i_*, calculated as follows:

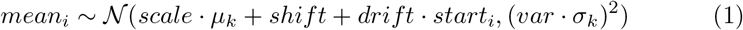

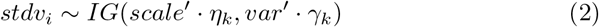

The most probable sequence is obtained through Viterbi decoding.

Performance of Nanocall at the moment of its publication, in March 0f 2016, was reportedly on par with that of 1D analysis by the Metrichor basecallers, or around 68% accuracy [7]. However, unlike the Metrichor basecallers, Nanocall does not support 2D analysis, which allowed for a much higher accuracy in the comparison by David *et al*. (around 85%) [7]. Since the Metrichor basecallers switched to a recurrent neural network-approach, its 1D accuracy proved significantly higher than that of Nanocall as well. Moreover, Nanocall is incapable of calling homopolymer stretches longer than its *k*-mer length, while current Metrichor basecallers are able to do so. Therefore, as the authors currently state on the Nanocall Github, it should be seen as a platform for testing out novel basecalling ideas, rather than a prime choice for basecalling MinION data [49].

#### DeepNano

Before Metrichor made its own switch from HMM- to RNN-basecalling, the open-source basecaller DeepNano [8] already implemented a form of RNN basecalling, booking a significant improvement in accuracy with respect to the then-current Metrichor version (corresponding to the SQK-MAP-006 kit, late 2015). DeepNano was written in Python, using the Theano library [50].

The RNN employed in DeepNano consists of 3 hidden layers of 100 units per layer for 1D basecalling and 4 hidden layers of 250 units for 2D. Rather than LSTM-nodes, as currently used in Metrichor basecallers, DeepNano implements gated recurrent units (GRUs) [51] to account for the vanishing gradient problem during parameter training (Figure 7B).

Many slight variations of the GRU exist, but all differ from regular LSTM-nodes in two aspects [45]. First, the GRU calculates the activation using only the activation of the previous step and the signal in the current step, omitting the usage of a separate memory input. Second, the GRU updates the hidden state using a single update gate, rather than separate forget and input gates. The design of GRUs is thus simpler than that of LSTMs, while performance assessments show that GRUs are able to achieve similar or slightly better accuracy in different learning tasks [45, 53].

DeepNano’s RNN was trained separately for (hairpin-connected) 1D base-calling and true 2D basecalling. As the exact alignment to the reference may be uncertain, the algorithm alternately trains the RNN weights for 100 iterations and then re-aligns the reads to the reference. At the moment of its release in March of 2016, DeepNano was reported by its authors to make more accurate calls than the Metrichor agent for both 1D calling (approximately 70% versus 77%) and 2D reads (approximately 88% versus 87%). However, as the Metrichor agent has since then switched to an RNN-approach as well these comparisons are outdated.

### 3.3 Assemblers

In the assembly stage, basecalled sequences are combined to reconstruct continuous parts of the genome as accurately and completely as possible. Assembly of nanopore reads requires a different approach than that of SGS reads; as nanopore reads are longer, finding a correct overlap should be easier, yet they are more error-prone, which increases the uncertainty of overlaps. Because of these differences a return of interest in overlap layout consensus (OLC) algorithms - which were at the peak of their popularity in the era of Sanger sequencing - is seen. De-Bruijn graph (DBG) assemblers, the more popular choice for SGS reads, were reported to return lower quality assemblies of nanopore reads than OLC-based methods, but proved faster in some cases [54]. A more detailed comparison between OLC and DBG assemblers, along with implementations of both approaches is discussed in this section. Software using traditional greedy extension algorithms (e.g. SSAKE) was found to perform decidedly less well in a *de novo* assembly setting, both in terms of assembly quality and required computational resources [54], and are therefore omitted here.

#### Theoretical effect of nanopore read properties on assembly

The Lander-Waterman model for sequence assembly can be used to obtain an insight into how long error-prone reads may affect the quality of the eventual assembly for OLC and DBG assemblers [55, 56]:

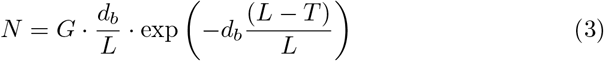

According to this model, the minimum number of contigs formed, *N*, is a function of genome size *G*, base coverage depth *d_b_* (i.e. the average number of times each base has been sequenced), read length *L* and a minimum overlap threshold *T*, chosen such that the selected overlaps are considered correct with a given confidence. For a genome of the same size, sequenced with the same coverage and assembled with the same overlap threshold, the number of contigs declines exponentially with increasing read length (Figure 8). On the other hand, the error-prone nature of nanopore reads will likely require a higher minimum overlap threshold for the same level of confidence, which causes an exponential rise in contig number. As DBG assemblers work with *k*-mers rather than individual bases, equation 3 does not directly apply in the same way. Introducing the k-mer coverage depth 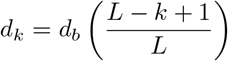 and considering that the minimum overlap between reads should be *k* (as this is required to establish a link between vertices), results in a slightly different form [56]^2^:

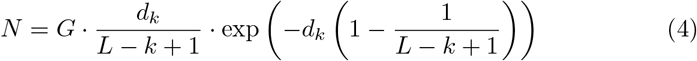

**Figure 8:**
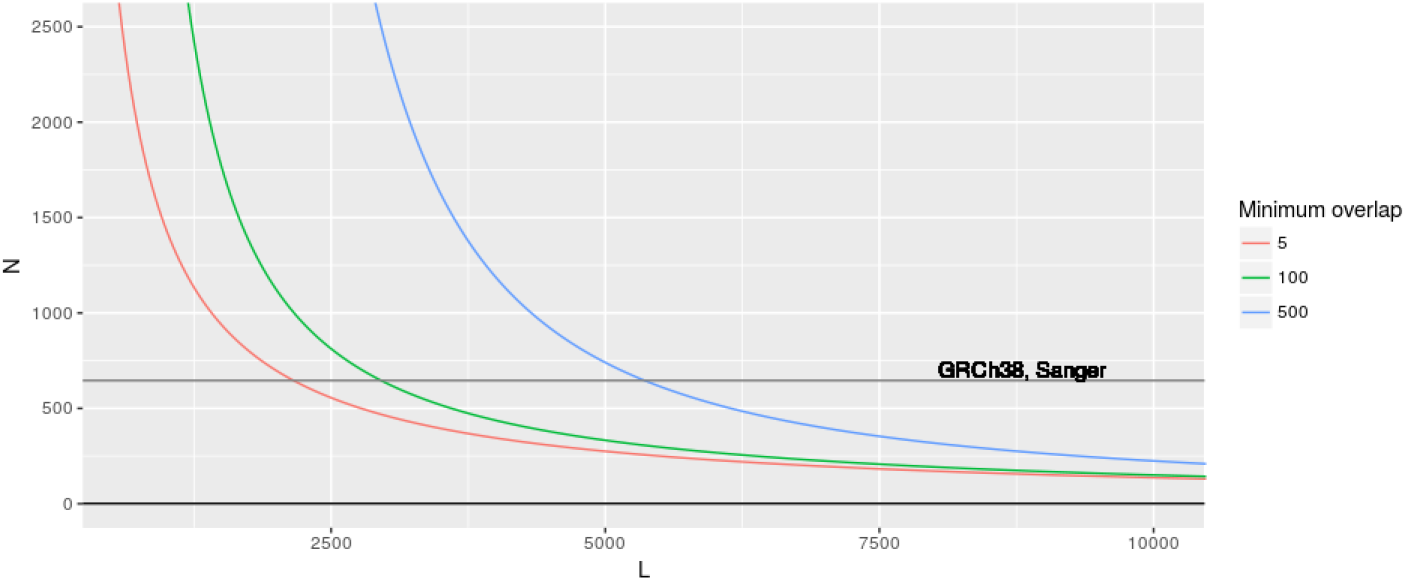
Theoretically expected number of contigs (N) when assembling a genome of 3 · 10^9^ bp (about the size of the human genome) with increasing read length (L) with error-free reads, according to the Lander-Waterman model [55]. As MinION read length for human reads currently averages at 10kb [4], an assembly in fewer contigs than the current GRCh38 genome [58] is possible according to this model, even with a considerable increase in overlap length, as would be required due to high basecalling error rate.

The implications for the effect of increased error rate and increased read length on the continuity remain the same; increased *k*-mer size to account for higher error rate and increased read length respectively cause an increase and decrease in the minimum number of contigs.

It should be noted that the Lander-Waterman model only offers an estimate of the potential minimal number of contigs, given ideal circumstances. It does not take the effect of remaining assembly errors on continuity into account, or the fraction of the genome consisting of repetitive elements. As discussed in the previous section the effect of errors can be severe, and arguably more so for DBG-assemblers than for OLC-assemblers. Below several assemblers that have been developed or adapted for use with nanopore data are discussed to give a more practical view of how the advantages and disadvantages of long error-prone reads pan out.

#### PBcR & Canu

Originally developed for the first human genome draft, the Celera assembly pipeline [59] and its extensions [36, 60, 61, i.a.] have remained a popular choice in a growing landscape of OLC assemblers. Briefly, the Celera assembler uses read overlaps to find contigs of which the structure can unambiguously be derived from overlap information, referred to as unitigs. It then separates repetitive unitigs from unique ones and attempts to orient the unique unitigs with respect to each-other. Where possible, gaps between unique unitigs are filled with repetitive elements. As a high read error rate is detrimental to the quality of the assembly [62], two different modifications to the pipeline are available. The PacBio corrected Reads (PBcR) algorithm, originally developed for the correction of PacBio reads suffering from similar error rates, uses accurate short reads mapped with high confidence to the long reads to correct errors. The assembly then proceeds as is usually done by Celera [63]. PBcR’s successor, Canu [36], provides a solution that is more integrated with Celera and does not require short accurate reads. The pipeline includes three stages; correction, trimming and assembly. Overlaps are found using the efficient Min-Hash alignment process (MHAP) [35], which hashes *k*-mers using different hash functions and for each hash function stores the smallest integer to which a k-mer of the sequence is hashed. Comparing the hashed *k*-mers per read results in initial overlap hits, which are then used to perform error correction by consensus seeking. By selecting overlaps for correction on quality but limiting the number of overlaps a read can contribute to, Canu attempts to prevent masking of true repeat variants. Shorter reads are used at this stage to improve accuracy of longer reads. In the trimming step, overlaps are recalculated to locate and filter out regions of low coverage and high error. Reads are overlapped two more times to correct specific types of errors (i.e. missed hairpin sections/adapters, chimeric reads) and adjust error rate per overlap, before the actual assembly phase starts. With adjustments to account for erroneous alignments and residual errors, assembly essentially follows the same procedure as CABOG, another Celera-based pipeline [60].

Due to its thorough yet relatively efficient correction steps, Canu is significantly more accurate than both its predecessor Celera/PBcR and Minimap/Miniasm [32]. In a benchmarking effort on *Enterobacter kobei* reads, it produced an assembly of higher contiguity, with fewer indels and mismatches. This is in line with the accuracy assessment by its authors [36].

#### Minimap & Miniasm

In terms of speed and computational efficiency, the OLC-based pipeline consisting of Minimap and Miniasm [39] has a definite advantage over other existing tools [32]. This efficiency was reached through the omission of the consensus step and the use of minimizers. Much like the Min-Hash algorithm used by Canu [36], a minimizer is a memory-efficient hashed representation of a sequence. Minimap computes the set of minimizers of a sequence, the “sketch”, by finding the *k*-mers represented by the smallest hash value within a certain window size of each position of the sequence (Figure 9). The inverse of each *k*-mer is also considered. Decreasing the window size will increase the returned number of minimizers and allow for more accurate alignment at the cost of increased computational requirements. Minimap then performs all-versus-all mapping by identifying hits between minimizers of different sequences. The found overlaps are passed on to Miniasm, which constructs an assembly graph. First, artefacts are trimmed from each read by removing all sequences outside the longest stretch with more than 3 times coverage. Then reads contained within other reads are removed and small bubbles, less than 50 kb in length, are popped (i.e. a consensus is taken in cases where paths split and later join up again). Finally, sequences can be extracted from stretches of the graph without multi-edges to form unitigs. The error rate at this point is practically the same as that in the raw reads, emphasizing that correct basecalling is essential for the eventual quality of the assembly. The graphical fragment assembly (GFA) output format of miniasm conveniently allows both graphing of the uncorrected assembly and addition of consensus error correction tools such as Nanopolish or Racon to the pipeline, though the latter increases walltime and computational cost severely [32].

**Figure 9:**
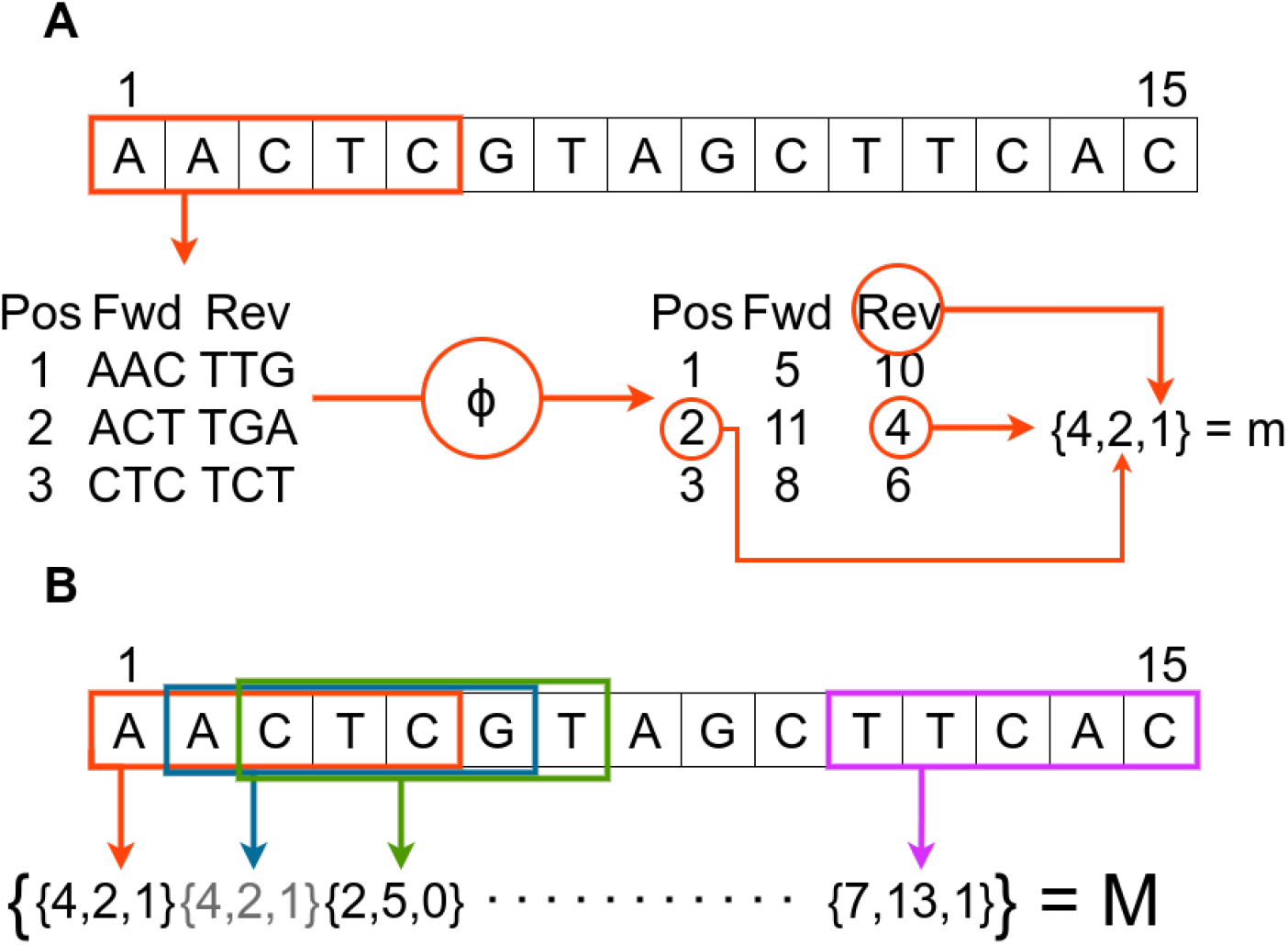
The minimizer sketch procedure, demonstrated on a 15-base read, using a window size of 5 and a *k*-mer value of 3. (A) To extract one minimizer m, all *k*-mers and their complements are listed and hashed to a distinct integer using hash function *ϕ*. The smallest hash value is then stored with its starting position and a binary value denoting the strand on which it was found (0 for forward and 1 for reverse). If a multiple of the smallest hash value is found within a window, all are stored. (B) The minimizers of all windows in the read are collected and stored as the sketch, M. As M is a set, double minimizers are removed.

In March of 2016, the authors of Minimap and Miniasm reported assembly of MinION reads of an *E. coli* genome in a single contig. In May of the same year Judge *et al*. assembled an *Enterobacter kobei* genome in 16 contigs with an N50 of 662 kb in two minutes, while the next fastest assembler (Canu) took two hours. However their benchmark showed that the omission of an error correction step caused the eventual assembly quality of *Enterobacter kobei* to be too low to properly assess by the QUAST analysis tool [32]. The inclusion of the Nanopolish consensus correction tool after Miniasm improved the quality of the assembly enough to perform a reliable quality analysis. However, the pipeline was still under-performing compared to other tools. As Nanopolish dramatically improves Miniasm assemblies primarily by correcting deletions in homopolymeric stretches [64], it is expected that better basecalling of homopolymer repeat regions beforehand would significantly improve Miniasm assemblies as well.

#### TULIP

As more reads are required to cover larger genomes and as the time required for all-vs-all overlapping increases quadratically with an increasing number of reads, it follows that the overlap step of OLC assemblers may take unfeasibly long for very large genomes. To tackle this issue, The Uncorrected Long read Integration Process (TULIP) takes a different approach to read overlapping [26]. Instead of all-vs-all alignment, short seed sequences are selected which the assembler then attempts to align with long reads. This drastically cuts down the overlapping complexity and makes efficient use of long reads to cover long stretches of the genome between the seed regions. The resulting graph represents seeds as vertices and the connecting reads as edges. In a graph cleaning step, vertices with multiple in- or outgoing edges are revisited. Spurious and superfluous edges are removed aggressively, thus producing a linear graph. Note that, as the name implies, TULIP does not perform basecalling error correction.

The success of assembly using TULIP highly depends on proper seed selection. To avoid spurious connections between reads, the seeds need to be sufficiently unique in the genome and contain few sequencing errors. If available, SGS reads may be used to construct seeds, although with the increasing accuracy of MinION reads the ends of long reads may be used as well. Apart from cutting out the need for SGS methods, the latter approach has the added advantage that pairs of seeds are connected by at least one long read. Furthermore, as TULIP is not able to assemble regions in which the gap between seeds is larger than the read length, a proper seed density over the entire genome is required. If a marker map is available for the genome, this information can be used to control the distribution of seeds in the selection process.

As a first demonstration of TULIP’s efficiency, Jansen *et al*. assembled the genome of the European eel *Anquilla anquilla* (approximately 850Mbp) with 18x coverage in three hours, requiring only 4.4GB of RAM and four threads [26]. The resulting assembly was more continuous than the SGS-based reference genome. As was the case with Minimap/Miniasm however, the current quality of MinION reads combined with the lack of an error correction step necessitates post-assembly correction. The authors further showed that missed seed alignments were the most commonly encountered issue during graph simplification, followed by tangled alignments due to repetitive seeds and spurious alignments. The seeds, constructed from short SGS reads, only underwent selection by uniqueness which did not lead to an equal distribution over the entire genome, however density remained high enough for successful assembly. The authors noted that assembly using the tips of MinION reads as seeds proved successful for *E. coli* genomes, but this has not been attempted for larger genomes yet (personal communication, May 1, 2017).

#### HINGE

Although long reads provide a definite edge when attempting to resolve repeat regions, issues may still occur if not all individual repeats are spanned by at least one whole read. In such cases, HINGE may provide a solution. Rather than attempting to resolve frayed rope structures in the assembly graph afterwards, HINGE pre-processes the reads to separate repeat regions that are entirely spanned by a read (and are thus more easily resolvable) from those that are not, and collapses the latter beforehand [37].

First, HINGE attempts to identify reads that wholly or partly overlap a repeat region. It does so by performing all-vs-all alignment and then selecting those reads of which a stretch aligns to a proportionally larger number of other reads than other parts. The intuition behind this is that reads from all copies of a repeat region existing in the genome align to each other, thus causing a characteristic abrupt increase in alignments for reads that overlap these repeat regions. Repeat regions covered entirely by at least one read can be easily resolved and are omitted from the following procedure. Of the reads lining the same repeat region, the reads that extend furthest into the repeat region (regardless of the location of the actual copy), are designated “hinges”. In the subsequent greedy extension of the hinges, the path will split at the hinge regions. Like Miniasm, HINGE outputs its assembly in the form of a graph. As its authors show, this is particularly useful for circular genomes.

HINGE provides an elegant solution to long repeat resolution, by separating resolvable regions from unresolvable ones on forehand. Its authors compared HINGE to Miniasm on PacBio reads of 997 circular bacterial genomes and found that overall, HINGE produced a completed genome in more cases than Miniasm could [37]. Whether the precaution taken by HINGE is necessary is dependent on the genome under consideration and the used reads; if the genome is known to contain repeats longer than most of the reads, the described approach would be justified.

#### Velvet

Velvet [65] is one of the older open-source DBG assemblers that are still maintained. It commences assembly by constructing two databases; one of unique *k*-mers per read and one of *k*-mers occurring multiple times in different reads. Each uninterrupted set of original *k*-mers per read is then merged, thus retaining more long-range information. Jumping from vertex to vertex using the information on the *k*-mers occurring multiple times, the edges are created. Their multiplicities are retained for error correction. Simplification is achieved by merging sequential nodes that do not branch. If short branches end in tips and are connected by an edge of lower multiplicity than other branches, they are removed. Bubbles are resolved by merging the paths if they are deemed similar enough. Lastly vertices that are covered by few reads at this stage are considered the product of chimeric reads and are removed.

Velvet’s performance on long error-prone reads has been compared to other assemblers on one occasion [54]. Although it performed better than the other DBG-assembler included in that benchmark (ABySS), it only occasionally provided a slight advantage in computational efficiency over OLC assembler PBcR/Celera and produced less contiguous assemblies.

#### ABruijn

While more traditional DBG assemblers performed relatively less well than OLC assemblers on assembling long error-prone reads, the adapted approach taken by the ABruijn assembler has shown more promise [38]. To account for the high error rate, ABruijn filters all *k*-mers occurring in the reads by their frequency; if a *k*-mer occurs relatively few times, it is assumed that it contains basecalling errors and it is removed. Then, *k*-mers are fused into so-called “solid strings”, sequences that contain no other occurring sequences as substring. The A-Bruijn graph is then drawn by representing solid strings as vertices and connecting them where connections exist in the reads. The edges are weighted by the number of positions between the first bases of the connected solid strings. The assembler consults the weights in this graph to quickly identify overlaps between reads, allowing to select on a minimum overlap length and maximum overhang length. The assembly graph is constructed by starting with the graph for an arbitrary read and continuously extending it by overlapping it with other reads. ABruijn also includes an error correction routine, during which a best consensus between reads is found by identifying low-error stretches and, in between those stretches, choosing the consensus sequence that maximizes the likelihood of the read sequences.

Unfortunately, the authors of ABruijn evaluated their assembler only on *E. coli* MinION reads of an older and more error-prone chemistry (R7.3), and compared their assembly to that of a Nanocorrect and Celera pipeline, which is currently considered obsolete. In this comparison ABruijn performs better, producing an error rate of 1.1% versus 1.5% for Nanocorrect/Celera, with the majority of errors being deletions. How ABruijn stacks up against currently popular assemblers optimized for MinION data on reads produced with up-to-date chemistries is unclear. The more extensive assessment on PacBio reads provided by the authors shows a dramatic decrease in error rate in te ABruijn assembly in comparison to Canu’s, but requiring comparable running time.

### 3.4 Post-assembly correction tools

A number of tools attempt to improve assemblies by remapping reads to the assembly and adapting the assembly to increase local resemblance to the reads. These tools may be essential to use after assemblers that do not include a consensus step themselves, such as Minimap/Miniasm and SMARTdeNovo, but have also frequently been used to polish assemblies produced by the other assemblers.

#### Nanopolish

Nanopolish attempts to find an optimal consensus between an assembly and the raw current signal output by the MinION, by iteratively proposing and evaluating small adaptations to the assembly based on the original reads [46]. The proposal mechanism for adaptations works in two steps. First, reads are aligned to the assembly and the resulting multiple alignment is divided in 50 bp subsequences of the assembly. For each read aligning partly or fully to a subsequence, sections in which 5-mers perfectly align to the assembly are detected. The consensus sequence between each pair of aligning sections is replaced by the aligned read subsequence, creating an initial set of alternative candidate sequences. In the second step, this set is further extended by proposing every possible one-base deletion, insertion and substitution in the previously generated candidate sequences. Of this set, the sequence maximizing the likelihood of observing the raw signal is picked. This process allows Nanopolish to explore a decent number of likely modifications, while remaining computationally tractable.

Polishing an assembly with Nanopolish was found to improve assembly quality, regardless of the assembly tool used. One study on *E. coli* sequencing data reported that identity to the reference genome rose from 89% to 99% when Nanopolish was applied after Minimap/Miniasm, while improvement after Canu was more modest (98.2% to 99.6%) [64]. Although it addresses all types of errors, a large part of the increase in accuracy is reached due to the correction of homopolymeric stretches. Despite its efficient searching heuristic of block replacement and mutation, running Nanopolish remains a time-consuming step; on the error-prone assembly produced by the otherwise quick Minimap/Miniasm pipeline, it may take up to a few extra days of CPU time to finish polishing [32, 64].

#### Racon

Racon corrects MinION assemblies by finding a consensus sequence between reads and the assembly through the construction of partial order alignment (POA) graphs. After alignment of the reads by a mapper of choice (e.g. Minimap or Graphmap), Racon segments the sequence and finds the best alignment between a POA graph of the reads and the assembly. By default, the alignment is performed using the Needleman-Wunsch algorithm, which can align sequence and POA graph with little adaptation (Figure 10). The alignment process is sped up by parallelization. Racon was reported by its authors to be two orders of magnitude faster than the popular (yet currently deprecated) Nanocorrect after assembly of an *E. coli* genome by Miniasm, albeit not quite as good at diminishing the error rate (to 1.31% versus 0.62% for Nanocorrect). Compared to consensus steps in Falcon [66] and Canu [36] on that same assembly, Racon remains an order of magnitude faster while producing similar error rates. A closer look at the error rate reveals that the majority of errors are indels, validating the need for homopolymer and tandem repeat corrections even in a pipeline containing a polishing step with Racon. Finally, the total genome size estimate following application of Racon was closer to the reference genome size than the estimates of Canu, Falcon and Nanocorrect.

**Figure 10:**
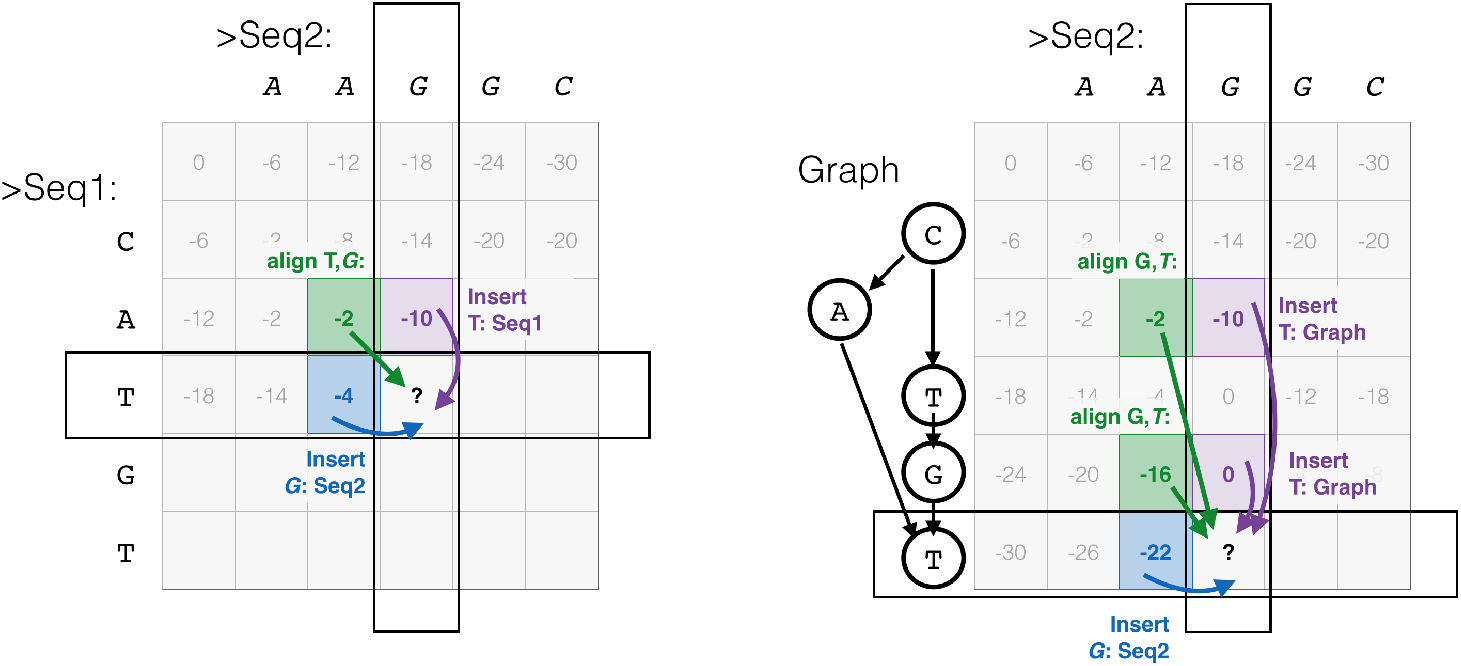
Needleman-Wunsch alignment of two sequences (left), compared to alginment of a partial order alignment (POA) graph and a sequence. In the latter case, the graph dictates which steps in the matrix are allowed. Taken from [67].

## 4 Assessing MinION basecalling software

In order to select the appropriate analysis tool from the mentioned options for a genome with a given set of characteristics, it is important that tools are rigorously benchmarked on several datasets bearing different characteristics. However, existing basecaller tools have not been evaluated as such. For some MinION-compatible assemblers, a benchmarking effort has been made [32, 54, 68], however these investigations were done using older and multiple MinION chemistries and basecaller tools, included relatively few assemblers and assessed performances mostly on bacterial genomes. Indeed, the image they sketch of assemblers may not provide today’s MinION users with the information required to make an informed choice for their user case.

Alternatively, users may want to perform their own performance analysis, using either their own data or the growing amount of freely available MinION data on the web [3, 69]. To aid such investigations, a recommendation of several appropriate metrics is given here, along with analysis tools that implement them where possible.

### Basecalling performance metrics

In general, an evaluation of the ability of a basecalling pipeline to make correct calls entails a comparison against a reference sequence. This reference may be acquired using Sanger or SGS methods and thorough curation. MinION Reads are mapped to the reference to obtain the sequence that is presumably underlying the read. It should be noted that the correctness of the reference sequence may not be guaranteed, yet it is assumed as the ground truth in basecalling performance assessments. The NanoOK benchmarking tool automates this process and outputs many of the metrics discussed below [70].

Plain basecalling accuracy, the fraction of correctly called bases in mapped reads, is an easy to understand metric and often used in basecaller quality comparisons [7, 8, 46]. A breakdown of misclassifications per type of error - insertion, deletion and substitution - and, in the case of a substitution, the involved base type, gives valuable added insight in any bias towards certain types of errors. The Phred-score [71] is a related often-reported quality metric for basecallers [72, 73] and is defined as −10log _10_(*Accuracy*). Correlation graphs between counts of *k*-mers, directly after basecalling or after assembly, and the reference may further characterize basecaller biases for calling certain *k*-mers [26, 46]. As a change in the basecalling routine is bound to have an effect on computing efficiency, the call time per base and memory should be reported as well.

### Assembler benchmarks

Drawing from Sanger and SGS assembly benchmarking efforts, a plethora of metrics is available for the evaluation of assembly quality. Many of these are implemented in QUAST [74], an often used assembler benchmarking tool [32, 46]. A selection of included metrics relevant in this work, i.e. those characterizing the size of constructed contigs and the correct-ness of the assemblies, is discussed here. In QUAST, a misassembly is defined as a point in a contig that is either mapped to a different chromosome, to the opposite strand or to a position of which the left end is more than 1 kb removed from the correct position in the reference. QUAST reports differences between the assembly and the reference in mismatches and indels per 100 kb. It should be noted that these are not the same metrics as those calculated in a basecalling performance benchmark; the used assembler or a post-assembly correction step may attempt to find a consensus between reads and as such diminish the error rate.

As noted previously, the long reads produced by the MinION are exceptionally suitable to construct larger, fewer contigs than possible using only SGS reads [32]. Thus, contig size and number are valuable metrics to assess this strength of MinION-based assemblies. QUAST reports these values and additionally provides several variants of the N*x* metric to further characterize contigs in relation to the assembly and genome size. More precisely, the N*x* value denotes the maximum contig length *L*, for which contigs of length larger than or equal to *L* cover *x* percent of all bases in the assembly [74]. The N50 value is most commonly reported. In a similar fashion, the NG50 denotes the maximum contig length for which contigs larger than or equal to this value cover half of the considered genome. As these scores characterize contig length without taking the actual quality of the assembly into account, Gurevich et al. [74] introduced the NA50 and NGA50 values, denoting the same metric calculated over contigs that are split at points of misassembly. Additionally QUAST can construct graphs showing how these N*x* values change with increasing minimum contig length.

QUAST also provides two metrics to characterize how the assembler covers the genome with the given reads, regardless of contig size. The genome fraction is defined as the number of bases covered by at least one mapped contig divided by the total length of the reference genome. The duplication ratio is defined as the total number of bases in aligned contigs divided by the number of bases covered in the reference. Thus, if many aligned contigs correspond to the same regions or overlaps are large, this value may be greater than 1.

Several additional metrics of interest were listed by Chu *et al*. [68]. Defining an overlap between reads as correct if found in a mapping of the reads to a reference, classic metrics of accuracy such as sensitivity, precision and false discovery rate (FDR) characterize how well an assembler was able to find correct overlaps, thus giving a lower-level indication of assembler performance. These metrics can be displayed in a receiver operator (ROC)-like curve, in which sensitivity is offset against FDR. Sensitivity and precision can also be combined into a single metric by taking their harmonic mean, which is commonly reported as the F1-score.

Lastly, an improvement in basecalling may have an effect on the computational efficiency of the assembler. Thus, it may prove insightful to compare CPU time and memory usage before and after a correction step in basecalling.

## 5 Discussion

Nanopore sequencing is a promising new venue in biology research. Inexpensive, small, capable of producing long reads and freed from the need for nucleotide labeling or amplification, it is conceivable that the MinION will make cost-effective, fast and portable *de novo* whole genome sequencing of even complex genomes possible in the future. In this review, an attempt was made to give an updated overview of the progress in this field, focusing in particular on *de novo* whole genome sequencing.

Available basecaller tools have been improving rapidly in accuracy. A notable recent improvement was made through the application of RNNs. For the next step in a typical sequencing routine, assembly, OLC-assemblers are currently considered the best option for accurate nanopore-based assembly. The choice of assembler should be adapted to the characteristics of the genome and the priorities of the user. Canu is a complete and accurate solution, while maintaining reasonable CPU times. However, for an assembler without included error correction, Minimap/Miniasm is able to produce a decent assembly and, without any post-assembly correction, is the fastest option available on smaller genomes. For large complex genomes, TULIP may be the more tractable alternative. Lastly, consensus error correction tools are currently highly useful. Notably, Racon provides a computationally more attractive alternative to the often-used Nanopolish.

Currently, the most prominent obstacle for *de novo* sequencing using the MinION is the high error rate of the reads. Improving basecalling accuracy would not only improve assembly quality in a direct manner, but may also allow more computationally efficient assembly.

The active research community surrounding the MinION has booked great progress in both the development of new applications and improvements on accuracy of existing ones. ONT also continuously works on improvements for both its hardware and software platforms, and regularly updates its users on this.

Although these updates often entail welcome new features or some form of accuracy improvement, it should be noted that this policy has also lead to some difficulties. Developers of tools may not be able to keep pace with ONT when evaluating, updating or calibrating their tools, and users may not always know which tool is suited best to their needs. As a result, most published studies, including tool benchmarking efforts, were conducted using older or multiple chemistries. Although such growing pains are to be expected for a novel fast-developing field of research, the MinION’s current state of development may allow for some increase in stability, thus giving the user community the time for proper evaluation.

## Footnotes

1 Estimate based on a purchase of 24 flowcells and a 1*D*/1*D*^2^ sequencing kit, 18th of May 2017

2 The form noted here is different from that noted in [56], as k is assumed to be the minimal overlap required to form an edge, while [56] assumes *k*+1. The interpretation given here is thought to be more in line with definitions of DBG methods in other publications (e.g. [57]).

## Authorship contribution statement

Carlos de Lannoy drafted the manuscript for this article and made included figures unless noted otherwise in the caption of figures. Judith Risse and Dick de Ridder regularly provided proofreading, feedback and suggestions for content.

## Acknowledgements

Giovanni Maglia provided helpful advice related to the physical basis of nanopore sequencing.

